# Urbanisation is associated with habitat-specific richness loss and community turnover across an ecotone

**DOI:** 10.1101/2025.09.13.675372

**Authors:** Alejandro Martinez, Sude Cakir, Marta García-Cobo, Diego Fontaneto, Michael Lemke, Maximilian Pichler, Nuria Sánchez, Thom Stoppelenburg, Jan-Berend Stuut, Willem Renema, Jan-Niklas Macher

## Abstract

Urbanisation can reduce biodiversity through environmental filtering, but its effects may differ among neighbouring habitats within ecotones. Beach–dune systems provide a useful test because strong environmental gradients occur over short distances and overlap with gradients in human pressure. We collected 660 sediment samples from 55 transects along the Dutch North Sea coast, covering established dunes, incipient dunes, the upper intertidal zone, and the lower intertidal zone. Using environmental DNA metabarcoding and habitat-specific joint species distribution models, we tested how distance-to-city was associated with community composition, operational taxonomic unit (OTU) richness, and OTU-level occurrence. We found compositional differentiation along the urban-proximity gradient in both dune habitats, particularly in incipient dunes, but not in the intertidal habitats. Across all habitats, communities far from cities were more similar to one another than communities near cities, indicating greater among-site compositional variation near urban areas. OTU richness and mean OTU occurrence probabilities increased with distance-to-city in the lower intertidal, upper intertidal, and established dunes. Incipient dunes showed no clear richness gradient or directional OTU-level response despite the strong compositional turnover, suggesting that the main response was community replacement rather than loss of OTUs. Overall, habitat position within the beach–dune ecotone influenced whether biodiversity responses to urban proximity led to directional filtering, compositional turnover, or increased community heterogeneity. Our findings show that biodiversity and impact assessments based on a single habitat may miss important ecological change in coastal habitats.

## Introduction

Human pressures often overlap with natural environmental gradients that shape ecological communities. The same pressure can therefore affect communities in neighbouring habitats in different ways. In some habitats, it may reduce richness, whereas in others it may alter taxonomic composition or increase community variation. These responses have different ecological implications, but they are often not distinguished when anthropogenic gradients are studied within one habitat type of an ecotone. Understanding the habitat-specific responses is important for predicting biodiversity change in landscapes where habitats are connected but environmentally different.

Beach–dune systems are useful for examining this issue because they span habitats that differ strongly in environmental conditions and exposure to human use. Communities in these ecotones change over short distances, from vegetated terrestrial dunes to sandy intertidal habitats. At the same time, the different habitats may be exposed to a shared gradient in urban proximity, which is associated with recreation pressure, beach management, infrastructure development, and other forms of human disturbance. Beach– dune ecosystems are also highly relevant for conservation because coastal zones are biologically diverse, provide ecosystem services such as coastal protection and recreation (McGranahan et al. 2007, Barbier et al. 2011), and more than 40% of the world’s population lives within 100 km of a shoreline (Seto et al. 2012, Neumann et al. 2015).

The pressures associated with shoreline development and recreation can act differently across the beach– dune gradient. Beach grooming removes wrack and organic matter that are essential to coastal food webs (Hyndes et al. 2022), infrastructure alters sediment dynamics, and recreational use can disturb habitats and species (Schlacher et al. 2008a,Bonte and Maes 2008, Martínez et al. 2020). The upper intertidal beach and the incipient dunes, i.e., the first line of young, emerging dunes, are directly accessible to beachgoers. These habitats are also naturally exposed to stress from desiccation and shifting sands. By contrast, the lower intertidal beach is partly buffered by frequent inundation, whereas established dunes are buffered by greater vegetation cover (Johnston et al. 2023) and are often less accessible. Therefore, the ecological effects associated with urban proximity potentially depend on habitat position within the ecotone.

Urban pressures can reduce the number of species, alter which species occur, and favour disturbance-tolerant generalists closer to cities (Schlacher and Thompson 2012, Bessa et al. 2014, Heerhartz et al. 2016, Peña-Alonso et al. 2019, Orlando et al. 2020, Torres et al. 2022). Such effects have been documented in different parts of beach–dune systems. Dune arthropod diversity can decline in small habitat patches (Van De Walle et al. 2025), trampling has a negative impact on arthropods (Comor et al. 2007, Bonte and Maes 2008), and urbanisation can reduce diversity in dunes (Chen et al. 2014, Abbate et al. 2019). In beach habitats, recreation, beach cleaning, driving, and artificial light can also affect animal communities (Fanini et al. 2014, Costa et al. 2020) (Gilburn 2012, Davies et al. 2016, Griffin et al. 2018) (Luarte et al. 2016). However, most studies focus on a single habitat within the beach–dune system. It therefore remains unclear whether neighbouring habitats respond similarly to the same urban gradient or whether their position along the ecotone changes the form of the biodiversity response. Urban proximity may reduce OTU richness in some habitats but primarily cause community replacement in others (Chase and Myers 2011, Cadotte and Tucker 2017). Previous work has compared community composition among beach and dune habitats with contrasting morphodynamics (Innocenti Degli et al. 2021), but tests of the same urban gradient across adjacent beach–dune habitats remain rare. Resolving these habitat-specific responses is also relevant for conservation and ecological monitoring under frameworks including the European Water Framework Directive (Ferreira et al. 2007) and the Habitats Directive (Delbosc et al. 2021).

A further limitation is that many studies focus on macroinvertebrates with body sizes greater than 1 mm. On heavily modified coasts such as the Dutch west coast, these taxa can be rare or absent (Janssen and Mulder 2005), limiting the applicability of traditional surveys that rely on macroinvertebrates. Smaller animals, including meiofauna with body sizes of approximately 40 μm to 1 mm, are abundant and respond rapidly to environmental change (Rodrı guez et al. 2003, Kotwicki et al. 2005, Martínez et al. 2025). They may therefore be useful for detecting responses to both natural environmental gradients and gradients in urbanisation (Pawlowski et al. 2018, Cordier et al. 2019)

Here, we combined environmental DNA metabarcoding (eDNA; (Taberlet et al. 2012) with scalable joint species distribution models (sjSDMs; (Pichler and Hartig 2021, Hartig et al. 2024) across the beach–dune gradient of the Dutch North Sea coast. We tested whether urban proximity was associated with richness loss or compositional turnover, or a combination of these responses, and whether the responses differed among lower intertidal, upper intertidal, incipient dune, and established dune habitats. Specifically, we tested two hypotheses:

**H1:** Urban proximity is associated with habitat-specific compositional turnover.

**H2:** Urban proximity is associated with habitat-specific filtering of OTU richness

We expected the strongest responses in the upper intertidal and incipient dunes because these habitats experience high natural stress, are easily accessible, and are often altered by infrastructure development. Figure 1 provides a conceptual overview of these predictions.

**Figure 1.**
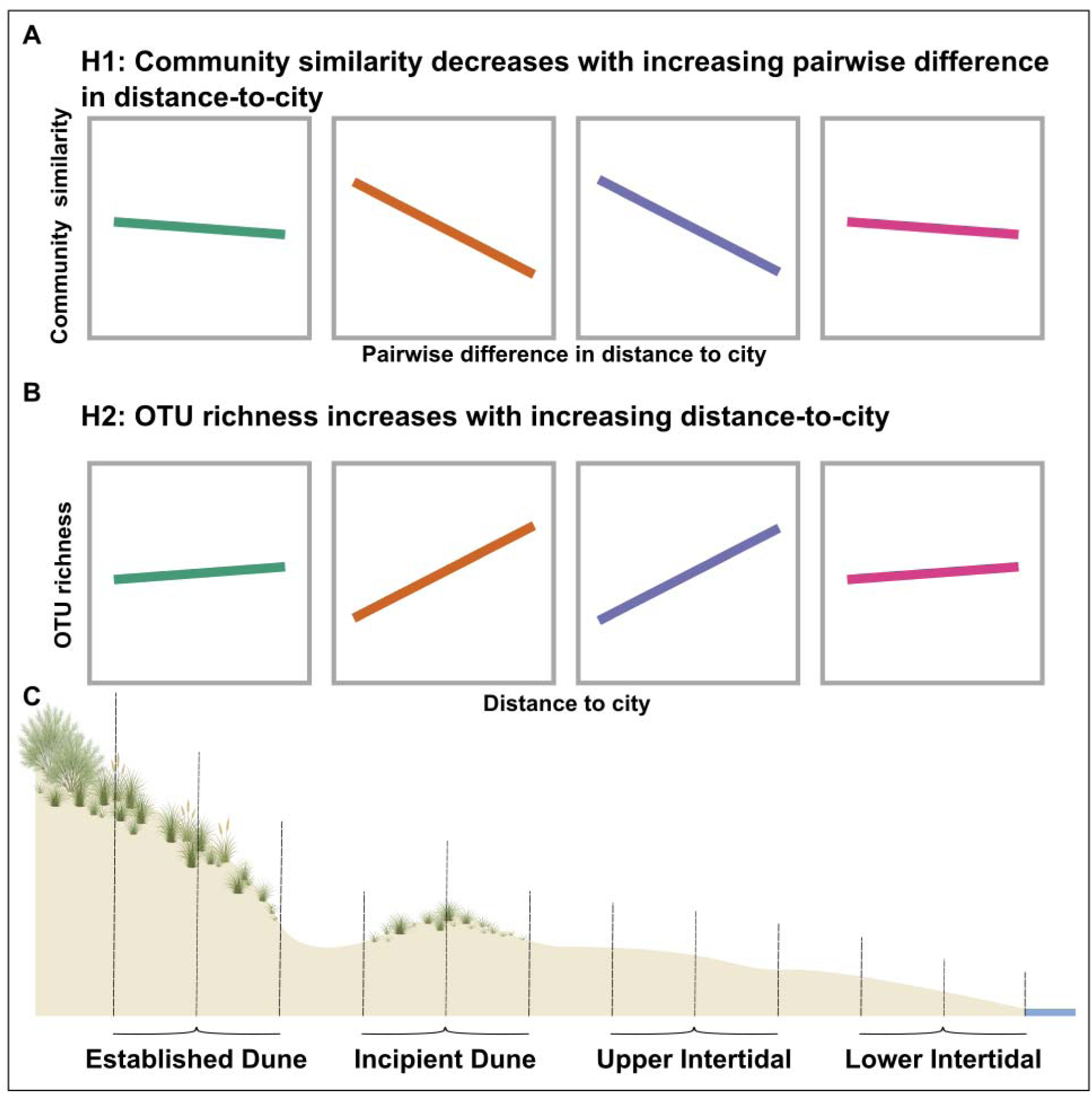
Conceptual overview of hypotheses and sampling design across the beach–dune gradient. Panels A and B show the predicted direction of urban-proximity effects within each habitat zone. A) H1: within-habitat community similarity is expected to decrease as pairs of sites become more different in distance-to-city, i.e. Sørensen similarity declines with increasing |Δ distance-to-city|. We expected this decline to be strongest in incipient dunes and the upper intertidal. B) H2: OTU richness is expected to increase with distance-to-city, i.e. richness is expected to be lower near cities, with the strongest increase in incipient dunes and the upper intertidal. C) The four sampled habitat zones are shown along a cross-shore beach–dune transect, from established dunes through incipient dunes to the upper and lower intertidal. Coloured lines in A and B indicate conceptual predictions for each habitat.

## Material and Methods

### Sampling

We collected 660 sediment samples from 55 transects along the Dutch coastline between Scheveningen and Zandvoort. The transects covered a gradient from urbanised beaches at Scheveningen, Katwijk, Noordwijk, and Zandvoort to less urbanised areas between these cities. Sampling locations are shown in Supplementary Figure S1, and coordinates are provided in Supplementary Table S1. Sampling was conducted between 25 July and 14 August 2023. At each transect, we sampled four zones of the beach– dune system (Figure 1c). In the established dunes, we collected three samples between the base and ridge of the dune, across the transition from grass cover to dune heath. In the incipient dunes, we collected three samples between the seaward edge of the first incipient-dune vegetation and the trough separating the incipient and established dunes. We also collected three samples from the upper half and three from the lower half of the intertidal zone. The upper and lower intertidal zones were defined as the upper and lower 50% of the total intertidal width, respectively. To minimise sampling bias due to edge effects, the outermost sampling positions were located approximately 10% of the habitat-zone width, and the samples within each zone were spaced equidistantly between these positions.

Sediment was collected with sterile 20-mL plastic cores inserted into the sediment to the 10-mL mark. The sampled sediment was then transferred to sterile 50-mL Falcon tubes. All samples were stored at −20°C until further processing. For each transect, we recorded urbanisation parameters using the Urbanisation Index developed for beaches (González et al. 2014). This included the density of buildings on the sand, machine cleaning of the beach, number of solid waste particles in the sand, intensity of vehicle traffic on the sand, and frequency of visitors. We also measured beach slope in degrees. Urban proximity was measured as distance-to-city, defined as the distance from each transect to the nearest main beach access point in the nearest city, in metres. We measured grain size separately for each habitat zone within each transect. For this, the three samples from each zone were pooled in equal volumes. Average grain size (in μm), was then measured using a LS13320 Particle Size Analyzer (Beckman-Coulter, USA).

### Molecular workflow

DNA was extracted from sediment samples using the SDE protocol (Bollmann-Giolai et al. 2020). We amplified a 313 base pair region of the mitochondrial cytochrome c oxidase I (COI) gene using LerayXT primers, which target a broad range of Eukaryota. Each PCR reaction contained 11.7 µl mQ water, 2 µl Qiagen CL buffer (10x; Qiagen, Hilden, Germany), 0.4 µl MgCl2 (25 mM; Qiagen), 0.8 µl Bovine Serum Albumin (BSA, 10 mg/ml), 0.4 µl dNTPs (2.5 mM), 0.2 µl Qiagen Taq (5U/µl), 1 µl of each Nextera-tailed primer (10 pMol/µl), and 2.5 µl of DNA template. PCR amplification started with an initial denaturation step at 96 °C for 3 minutes. This was followed by 30 cycles of denaturation at 96 °C for 15 seconds, annealing at 50 °C for 30 seconds, and extension at 72 °C for 40 seconds. The PCR ended with a final extension step at 72 °C for 5 minutes. We processed six negative controls together with the samples to check for contamination. These controls contained Milli-Q water instead of DNA template (Milli-Q, Merck, Kenilworth, USA). After the first PCR, the samples were cleaned using AMPure beads (Beckman Coulter, Brea, United States) at a 0.9:1 ratio to remove short fragments and primer dimers. A second PCR with individually tagged primers followed the same protocol, using the PCR product from the first PCR as template, but reducing cycle number to 10. DNA concentrations were measured using the FragmentAnalyzer (Agilent Technologies, Santa Clara, CA, USA) with the DNF-910 dsDNA kit (35–1500 bp). Samples were pooled equimolarly, and the final library was cleaned with AMPure beads and sequenced on two Illumina NovaSeq 6000 runs at Baseclear, Leiden, The Netherlands.

### Bioinformatic processing of community metabarcoding data

Raw metabarcoding reads were processed using APSCALE (v.1.6.3) (Buchner et al. 2022) We used a maximum percentage difference of 10%, a minimum overlap of 50 bp, and a minimum sequence length of 300 bp. All other settings, including LULU filtering, were kept at their default values. Sequences were clustered into Operational Taxonomic Units (OTUs) at 97% sequence similarity. OTUs detected in the negative controls were removed using a subtraction approach. For each OTU found in the negative controls, the total number of reads detected in the controls was subtracted from the read count of that OTU in all other samples (Macher et al. 2023). Samples with less than 5% of the median read count across all samples were removed from further analyses. We also removed OTUs with less than 0.01% read abundance per sample, to reduce the influence of low-level tag jumping (Schnell et al. 2015). Taxonomic assignment was done using the NCBI GenBank database. This database was supplemented with meiofauna COI barcodes generated during associated taxonomic workshops (Macher et al. 2024). OTUs that could not be assigned to an animal reference sequence with at least 90% similarity were excluded from further analysis.

### Urbanisation gradients and covariates

We used distance-to-city as the main measure of urban influence. This was measured as the distance from each transect to the main beach access point of the nearest city. To test whether distance-to-city captured the main urbanisation gradient in the study area, we correlated it with the disturbance indicators recorded for each transect. These indicators were buildings on sand, beach cleaning, vehicle traffic, and visitor frequency. Beach waste showed no variation across the sampling area and was therefore not analysed further. We used Spearman rank correlations for these tests. Distance-to-city was strongly correlated with all disturbance indicators (ρ = −0.60 to −0.85; all BH-adjusted p ≤ 5.1 × 10⁻ ). This showed that urban disturbance increased closer to cities, and we therefore used distance-to-city as the main urban predictor in all models. As additional environmental predictors, we used sediment grain size in μm and beach slope in degrees. We also included latitude and longitude of each transect to account for spatial patterns along the coastline. All continuous predictors were centred and scaled before modelling.

### Joint Species Distribution Models (jSDM)

We used habitat-specific joint species distribution models to test how urban proximity was related to community composition. Models were fitted with the R package sjSDM, which allowed us to model shared responses to environmental predictors, spatial structure, and residual co-occurrence among OTUs. We fitted separate models for the lower intertidal, upper intertidal, incipient dunes, and established dunes.

Before modelling, the three cores per habitat zone and transect were pooled by summing read counts. Within each habitat, we modelled OTU presence-absence. We only included OTUs found in at least five transects within that habitat, because jSDMs can be unstable when many taxa are rare (Pichler and Hartig 2021, Cai et al. 2025). For each habitat, OTU presence-absence was modelled with a binomial probit model. The environmental predictors were distance-to-city, sediment grain size, and beach slope. To account for spatial structure along the coast, we included latitude and longitude as a flexible smooth surface by adding their linear terms, squared terms, and their interaction. We also included a regularised biotic association structure to model residual co-occurrence among OTUs after accounting for the environmental and spatial predictors. To reduce overfitting, we used weak regularisation for the environmental and spatial terms (λ = 0.001) and for the biotic association structure (λ = 0.0001) (Pichler et al. 2025). Models were fitted for 1,000 iterations using the default optimiser settings and a learning-rate scheduler with a reduce factor of 0.9.

We treated the habitat-specific jSDMs as the primary inferential framework. Community-level quantities that are not direct model parameters, including predicted OTU richness, expected Sørensen similarity, and near–far similarity contrasts, were therefore summarised from simulations of the fitted models conditional on the observed site covariates.

### Community similarity along the urban gradient

For each habitat, we used the fitted jSDM to simulate 300 presence-absence communities for each site. For each simulation, we calculated Sørensen similarity for every pair of sites. We then averaged these values across simulations to obtain one expected similarity matrix for each habitat. We summarised whether model-predicted community composition changed along the urban-proximity gradient. For this, we related mean Sørensen similarity to the absolute difference in distance-to-city between each pair of sites, written as |Δ distance-to-city|. We used a linear model to estimate the direction and strength of this relationship. A negative slope means that communities become less similar when sites are farther apart along the urban proximity gradient. Because each transect occurs in several pairwise comparisons, we did not use the linear model alone for inference. Instead, we used a permutation test. We used 999 permutations to generate the null distribution. In each permutation, distance-to-city values were shuffled among sites, |Δ distance-to-city| was recalculated for all site pairs, and the slope was refitted. We then compared the model-derived slope to this null distribution. We interpreted an urban-gradient relationship as supported only when the slope was unlikely under this permutation approach.

### Near-far comparisons from jSDM simulations

As an additional analysis, we used the jSDM simulations to compare community consistency near and far from cities. Within each habitat, we defined near sites as those in the lowest quartile of distance-to-city, meaning ≤ 25th percentile. Far sites were defined as those in the highest quartile, meaning ≥ 75th percentile. Sites in the middle 50% of the gradient were not used in these contrasts. For each habitat and each simulation draw (nsim = 300), we calculated mean Sørensen similarity for three types of site pairs: near–near pairs, far–far pairs, and near–far pairs. To summarise differences in community consistency between the near and far ends of the gradient, we calculated contrasts within each simulation draw, for example far–far similarity minus near–far similarity, or near–near similarity minus far–far similarity. We then summarised these contrast distributions across draws. We summarised the three pair categories using their means and 95% intervals across simulation draws. Pairwise contrasts were calculated within each draw, and statistical support was summarised using Benjamini–Hochberg-adjusted simulation p-values.

### OTU richness along the urban gradient

We estimated local diversity from the fitted community models using the jSDM simulations. For each simulation draw, we counted the number of OTUs predicted to be present at each site. Site-level richness was then summarised as the mean and 95% interval across the 300 simulation draws. To describe how richness changed along the urban gradient, we fitted a linear model between mean model-estimated richness and distance-to-city.

### OTU-level response to urban proximity

As a complementary analysis, we summarised OTU-level responses to urban proximity by extracting the jSDM coefficient (β) and standard error for the distance-to-city predictor for each OTU in each habitat. We then used a random-effects meta-analysis with REML to estimate the mean response across all OTUs within each habitat. We also repeated this analysis for OTUs assigned to phyla represented by at least five OTUs in that habitat. This analysis was used to summarise the overall direction of OTU occurrence responses, while accounting for uncertainty in the coefficients of individual OTUs. We classified group-level responses as positive when the 95% confidence interval of the mean effect was entirely above zero. This indicated higher occurrences farther from cities. We classified responses as negative when the interval was entirely below zero, indicating higher occurrence closer to cities.

## Results

### Urban-gradient structuring of community similarity

After bioinformatic processing, we retained 358 animal OTUs (see Supplementary Table S2 for the full read table and taxonomic annotations). We fitted separate habitat-specific jSDMs to test how urban proximity related to community composition within each habitat. Model performance differed among habitats, with explained variation highest in the incipient dunes (R² = 0.45) and established dunes (R² = 0.41), intermediate in the upper intertidal (R² = 0.36), and lowest in the lower intertidal (R² = 0.26).

We first assessed whether inferred community similarity changed continuously along the urban proximity gradient, by testing whether Sørensen similarity declined as pairs of transects became more different in distance-to-city (|Δ distance-to-city|). This relationship was habitat-specific and strongest in the dune habitats (Fig. 2; Supplementary Table S3). Because pairwise similarities are non-independent, we interpreted support for these relationships using permutation tests rather than the p-values of the fitted linear trends. The clearest signal was found in the incipient dunes (slope = −1.28×10⁻ , p perm = 0.001), followed by the established dunes (slope = −5.94×10⁻ , p perm = 0.033), indicating increasing compositional differentiation with separation along the urban-proximity gradient. Although the fitted slopes were also negative in the upper and lower intertidal, these relationships were not supported by permutation tests (upper intertidal: p perm =0.567; lower intertidal: p perm =0.402), so we did not interpret them as evidence for continuous urban-gradient structuring in the intertidal habitats.

**Figure 2:**
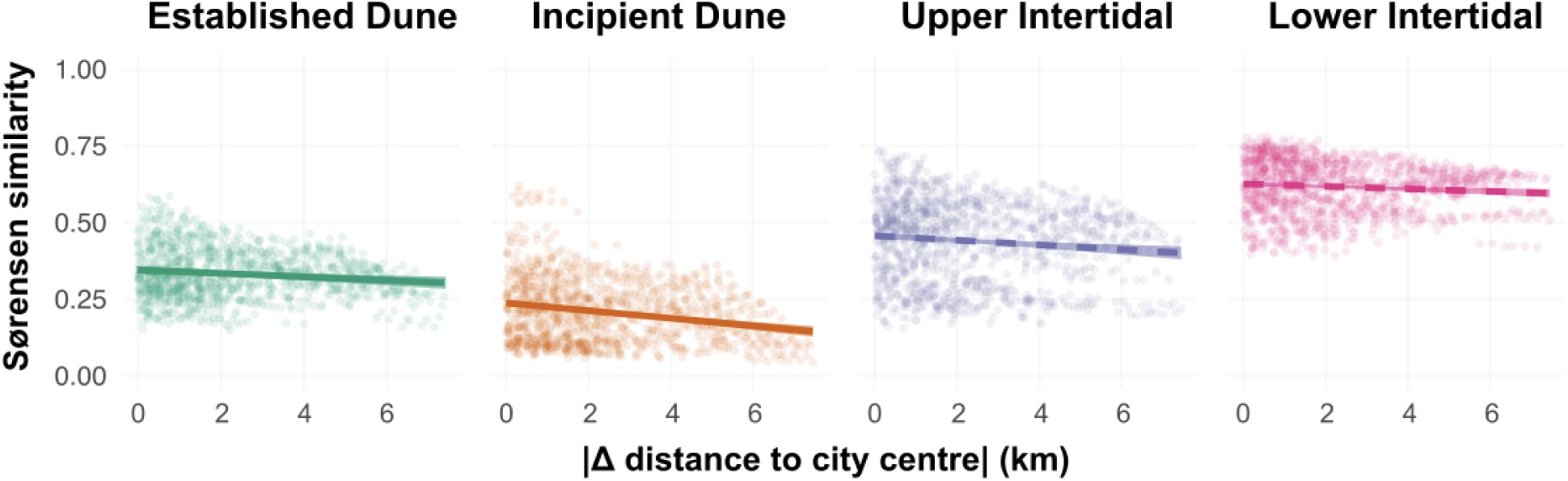
Urban proximity and community composition differentiation within habitats. Relationship between mean Sørensen similarity and pairwise separation of sampling sites in urban proximity (|Δ distance-to-city|, shown in km), shown separately for each habitat zone. Points show pairwise similarities; the line shows the fitted linear trend with a descriptive 95% confidence ribbon. Solid lines indicate permutation support (p < 0.05); dashed lines indicate no permutation support.

### Near–far contrasts show greater community consistency far from cities

The gradient analysis tested whether community similarity changed gradually with increasing separation in distance-to-city. Because pairs with low |Δ distance-to-city| can include both near–near and far–far comparisons, it does not show whether communities are equally consistent at both ends of the urban gradient. We therefore compared communities at the near and far ends of the gradient.

Across habitats, community similarity was highest among pairs where both sites were far from cities (Fig. 3; Supplementary Table S4). In every habitat, similarity between two far-from-city sites was higher than similarity for mixed near–far pairs (all BH-adjusted p ≤ 0.010), indicating that far-from-city communities were compositionally more consistent and distinct from near-city communities. Far–far similarity was also higher than near–near similarity in all habitats, indicating that near-city communities were more variable in composition. Mean within-far similarity ranged from 0.383 in the incipient dunes to 0.671 in the lower intertidal, whereas within-near similarity ranged from 0.199 to 0.606 and near–far similarity ranged from 0.124 to 0.590. These contrasts describe differences between the ends of the urban gradient and should not be interpreted as evidence for a continuous gradient response in all habitats.

**Figure 3:**
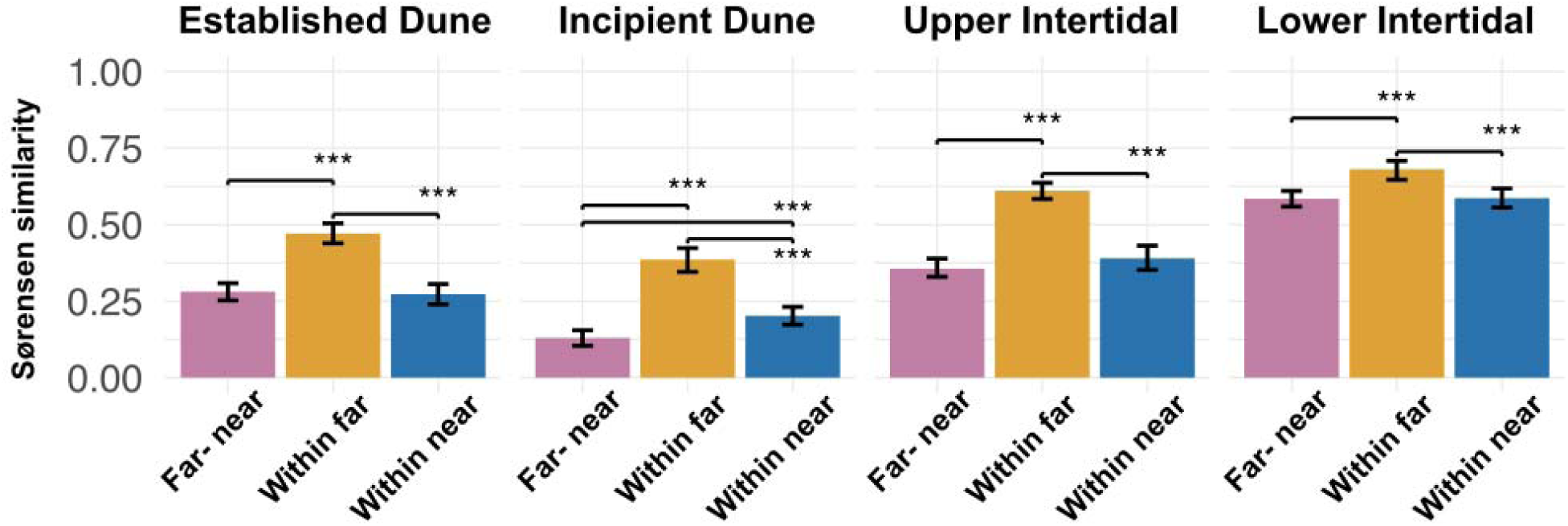
Near-far patterns in community similarity within habitats. Mean Sørensen similarity is shown for three categories within each habitat zone: within near (both sites near a city), within far (both sites far from a city), and near-far pairs (one near, one far). Black error bars indicate the mean and 95% interval across draws (n=300). Statistical support for contrasts among categories was assessed using simulation-based paired contrasts with Benjamini-Hochberg correction. Summary statistics and adjusted pairwise comparisons are provided in Supplementary Table S4.

### OTU richness along the urban gradient

Across habitats, model-estimated OTU richness generally increased with distance-to-city, although the strength of this relationship differed between habitats (Fig. 4; Supplementary Table S5). The clearest richness gradient was found in the upper intertidal (slope = 1.810, p < 0.001, R² = 0.562). Richness also increased with distance-to-city in the lower intertidal (slope = 0.819, p < 0.001, R² = 0.339) and established dunes (slope = 1.241, p = 0.005, R² = 0.165). In contrast, the incipient dunes showed no clear richness trend (slope = 0.006, p = 0.990, R² < 0.001). Thus, richness patterns suggest a directional filtering response in most habitats, but not in the incipient dunes.

**Figure 4:**
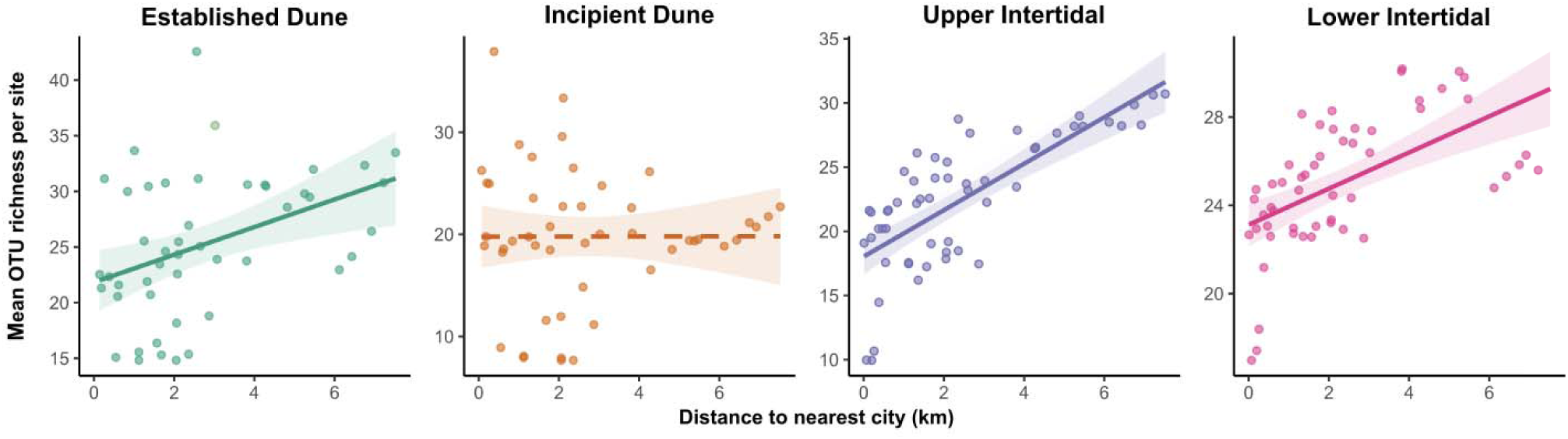
OTU richness along the urban proximity gradient. Relationship between distance-to-city (km) and OTU richness per site, shown separately for each habitat. Points show site-level mean richness across simulation draws (nsim = 300). Lines show fitted linear trends (mean richness ∼ distance-to-city) with 95% confidence ribbons. Line type indicates statistical support from the linear model (solid: p < 0.05; dashed: p ≥ 0.05). Full slope estimates are provided in Supplementary Table S5.

### Taxon-level responses to urban proximity

Mean OTU-level jSDM coefficients showed generally positive responses to distance-to-city, indicating that OTU occurrence tended to be higher farther from cities in most habitats (Fig. 5; Supplementary Table S6). Across all OTUs, mean responses were positive in the upper intertidal (β = 0.306, p < 0.001), lower intertidal (β = 0.200, p < 0.001), and established dunes (β = 0.180, p < 0.001). In contrast, OTUs in the incipient dunes showed no consistent directional response (β = 0.057, p = 0.214).

**Figure 5.**
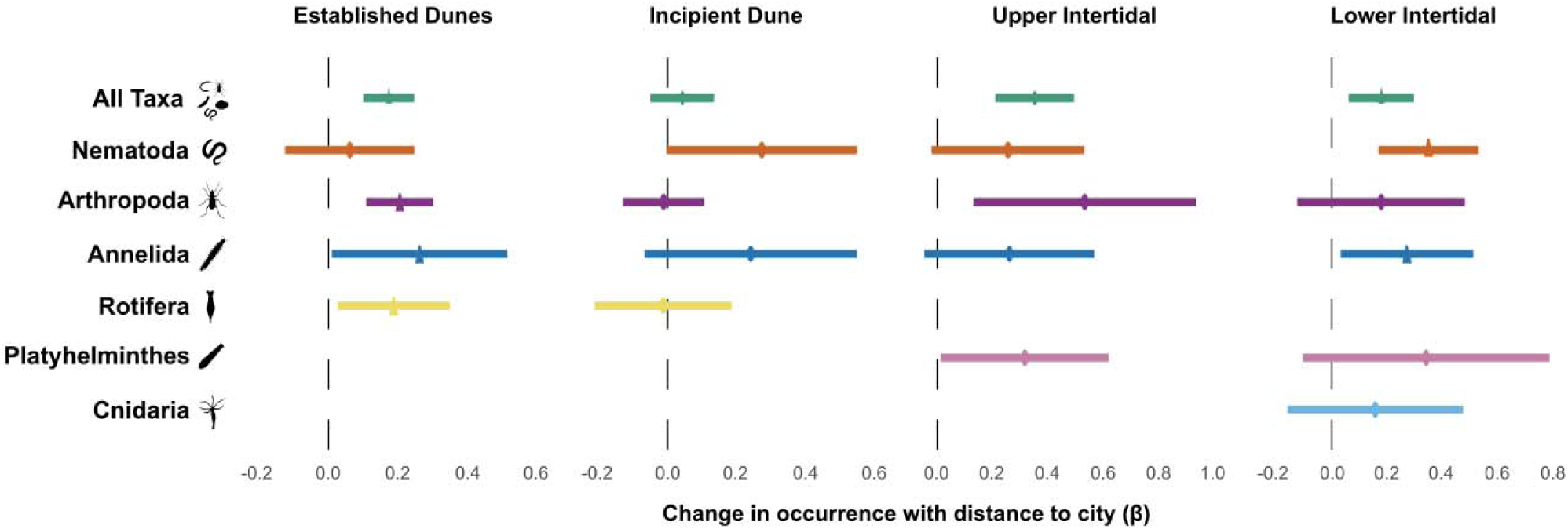
Directional responses of OTUs to urban proximity across habitats. Horizontal bars show the mean jSDM coefficients (β) ± 95% confidence intervals for the effect of scaled distance-to-city on OTU occurrence within each habitat. Positive values indicate higher occurrence probabilities at sites farther from cities, negative values indicate higher occurrence closer to cities, and confidence intervals overlapping zero indicate no consistent directional response. Estimates are shown for all OTUs and for OTUs assigned to common phyla. Full coefficient summaries are provided in Supplementary Table S6.

Phylum-level patterns were broadly consistent with the whole-community results but differed among habitats. In the established dunes, Annelida, Arthropoda, and Rotifera showed supported positive mean responses. In the upper intertidal, all analysed phyla had positive point estimates, but only Arthropoda and Platyhelminthes showed supported positive responses. In the lower intertidal, positive responses were supported for Annelida and Nematoda. In the incipient dunes, none of the analysed phyla showed a supported directional response. No phylum showed a supported negative mean response.

## Discussion

Human pressures often overlap with natural environmental gradients, yet neighbouring habitats may respond to the same pressure in different ways. Our results show that urban proximity was associated with distinct forms of biodiversity change across the beach–dune ecotone, and provided partial support for bothhypotheses.

Compositional turnover along the urban gradient was supported in the dune habitats, particularly in the incipient dunes, but not in the intertidal habitats (H1). By contrast, OTU richness indicated directional filtering in the lower intertidal, upper intertidal, and established dunes, but not in the incipient dunes (H2).

The complementary OTU-level analysis showed similar directional occurrence responses in these three habitats. In addition, comparisons between communities near and far from cities showed that near-city communities were more variable in composition than those farther away. Together, these findings show that habitat position within an ecotone can influence whether biodiversity responses to urban proximity are expressed primarily as richness loss, compositional turnover, or increased community heterogeneity.

### Urban proximity is linked to stronger community turnover in dunes

We found the clearest relationship between urban proximity and community composition in the dune habitats. In both dune zones, community similarity declined as transect pairs became more different in distance-to-city, whereas neither intertidal habitat showed such a relationship. Habitat position within the ecotone may therefore influence whether compositional change is structured continuously along the urban-proximity gradient.

This relationship was strongest in the incipient dunes. As early-successional habitats that are directly accessible to beachgoers, the incipient dunes may be more strongly exposed to changes associated with trampling, beach management, and infrastructure development. These pressures may influence habitat stability, organic input, or microclimate and thereby affect which taxa persist. Transitional habitats may be especially responsive to such environmental changes (McDonnell and Pickett 1990, Griffin et al. 2018). Because incipient dunes also contribute to dune development, changes in their animal communities may have implications for ecosystem functioning and coastal protection (McGuirk et al. 2022).

Established dunes also showed compositional differentiation along the urban-proximity gradient, but the mechanisms underlying this process may differ from those in incipient dunes. Established dunes are more vegetated and less exposed to hydrodynamic disturbance and direct disturbance like trampling. Urbanisation may instead affect these habitats through changes in vegetation structure, artificial lighting, and fragmentation by beach-access paths (Purvis et al. 2015, Aguilera et al. 2022). Such changes can change habitat conditions and favour disturbance-tolerant taxa (McKinney 2006, Bonte and Maes 2008).

The absence of supported turnover in the intertidal habitats may reflect the stronger influence of hydrodynamic processes and short-term environmental variability in these habitats. Frequent inundation and sediment movement may weaken or obscure patterns of composition turnover associated with urban proximity. However, previous studies have shown that urbanisation and recreation can affect sandy-beach macroinvertebrates (Gilburn 2012, Schooler et al. 2019). Our results therefore do not prove that urban pressures are not important in intertidal habitats. Rather, the effects of urbanisation may be more difficult to detect using sediment eDNA without direct measurements of pressures such as trampling and beach-grooming frequency. Furthermore, temporal replication may be needed to separate the effects of urban proximity from natural environmental variability (Defeo and McLachlan 2005, Schlacher et al. 2008b).

### Near-city communities are more heterogeneous

The near–far analysis showed a different component of community structure. While the continuous-gradient analysis tested whether communities became less similar as sites became more separated in distance-to-city, the near–far contrasts tested whether communities at either end of the gradient differed in their internal consistency. We found that across all habitats, far-from-city communities were more similar to one another than near-city communities, indicating greater among-site compositional variation close to cities.

This pattern differs from the common expectation that urbanisation homogenises ecological communities (McKinney 2006, Schooler et al. 2019). However, homogenisation can depend on the spatial scale and habitats being compared. One possible explanation is that local pressures and habitat conditions vary more among sites near cities, whereas sites farther from cities may experience more similar habitat conditions or more consistently low levels of disturbance. This interpretation is consistent with studies showing that human activities can increase variation in community composition in sites close to or in urban areas (Magura et al. 2018).

Differences among categories were strong in the dune habitats and upper intertidal but comparatively smaller in the lower intertidal In the incipient dunes, near-city communities were also more similar to one another than to far-from-city communities, suggesting clear compositional separation between the two ends of the gradient. In the lower intertidal, differences in community consistency were comparatively smaller, possibly because hydrodynamic processes strongly influence community composition (Defeo and McLachlan 2005, Harris et al. 2011), which may obscure the associations with urban proximity.

### Urban proximity is consistent with filtering in most habitats

Richness and OTU-level occurrence patterns were consistent with directional filtering in the lower intertidal, upper intertidal, and established dunes. In these habitats, OTUs were on average more likely to occur farther from cities, and the estimated richness generally increased with distance-to-city. These patterns suggest that conditions associated with proximity to cities may exclude some taxa or reduce their probability of occurrence, as expected when urbanisation effects act as an environmental filter (McKinney 2008, Martínez et al. 2020, Corte et al. 2022). The upper intertidal showed the clearest richness and whole-community occurrence responses. The mean coefficients were positive for all analysed phyla, but supported positive responses were found only for Arthropoda and Platyhelminthes.

Incipient dunes showed no clear richness gradient and also no consistent directional OTU-level response. The primary response in this habitat therefore appears to be replacement among OTUs rather than a reduction in OTU number. Communities near cities may represent compositionally different rather than simply species-poor versions of communities farther away from cities. Similar combinations of compositional change without richness loss have been reported in coastal plant communities (Del Vecchio et al. 2016) and other ecosystems (Lehnert et al. 2013, Hillebrand et al. 2018).

### Implications and future directions

Our findings show that biodiversity responses to an anthropogenic gradient cannot necessarily be inferred from one habitat or one biodiversity metric. Neighbouring habitats differed in whether their clearest response to urban proximity involved directional changes in richness and occurrence, compositional turnover, or greater among-site variation. Monitoring based only on richness may therefore overlook changes in community structure, particularly in transitional habitats.

Further work is needed to identify the mechanisms underlying these habitat-specific associations. The distance-to-city captured the main urban gradient in out study area, but it is a proxy and dis not allow us to directly determine which pressures connected to urban development caused the observed patterns. Direct measurements and manipulative experiments on the effects of trampling, beach grooming, artificial light, and nourishment could help separate the effects of individual pressures (Dugan et al. 2003, Schlacher et al. 2016). Although eDNA metabarcoding allowed us to include small and often overlooked meiofauna in the study of urbanisation impacts, the taxonomic resolution of these organisms depends strongly on the completeness of molecular and taxonomic reference databases, which need to be built for more regions (Castro et al. 2021, Macher et al. 2024). Applying the same framework to other coastlines would help determine whether the patterns observed in our study are general features of beach–dune ecotones or are strongly influenced by local management and environmental conditions. Furthermore, sampling adjacent shallow-subtidal habitats would broaden the ecotone perspective because these habitats may also be affected by coastal development (Corte et al. 2022).

## Conclusions

The effects associated with urban proximity differed among adjacent habitats within the beach–dune ecotone. Compositional replacement dominated in incipient dunes, whereas the other habitats showed patterns consistent with directional filtering. The greater compositional variation among near-city communities further showed that urban influence can affect the consistency of ecological communities. Predicting and monitoring biodiversity change across ecotones therefore requires comparisons among connected habitats and consideration of multiple dimensions of community structure.

